# Differential effects of PTH (1-34), PTHrP (1-36) and abaloparatide on the murine osteoblast transcriptome

**DOI:** 10.1101/2023.01.11.523646

**Authors:** Michael J Mosca, Zhiming He, Florante R. Ricarte, Carole Le Henaff, Nicola C. Partridge

**Affiliations:** Department of Molecular Pathobiology, New York University College of Dentistry, New York, NY, 10010, USA; Vilcek Institute of Graduate Biomedical Sciences, New York University School of Medicine, New York, NY, 10016, USA

**Keywords:** Osteoblast, Parathyroid hormone (PTH), Osteoporosis, Abaloparatide, RNA-Seq, Signal Transduction

## Abstract

Teriparatide (PTH(1-34)) and its analogs, PTHrP(1-36) and abaloparatide (ABL) have been used for the treatment of osteoporosis, but their efficacy over long-term use is significantly limited. The 3 peptides exert time- and dose-dependent differential responses in osteoblasts, leading us to hypothesize that they may also differentially modulate the osteoblast transcriptome. We show that treatment of mouse calvarial osteoblasts with 1 nM of the 3 peptides for 4 h results in RNA-Seq data with PTH(1-34) regulating 367 genes, including 194 unique genes; PTHrP(1-36) regulating 117 genes, including 15 unique genes; and ABL regulating 179 genes, including 20 unique genes. There were 83 genes shared among all 3 peptides. Gene ontology analyses showed differences in Wnt signaling, cAMP-mediated signaling, bone mineralization, morphogenesis of a branching structure in biological processes; receptor ligand activity, transcription factor activity, cytokine receptor/binding activity and many other actions in molecular functions. The 3 peptides increased *Vdr, Cited1* and *Pde10a* mRNAs in a pattern similar to *Rankl*, i.e., PTH(1-34) > ABL > PTHrP(1-36). mRNA abundance of other genes based on gene/pathway analyses, including *Wnt4, Wnt7, Wnt11, Sfrp4, Dkk1, Kcnk10, Hdac4, Epha3, Tcf7, Crem, Fzd5, Pp2r2a*, and *Dvl3* showed that some genes were regulated similarly by all 3 peptides; others were not. Finally, siRNA knockdowns of SIK1/2/3 and CRTC1/2/3 in PTH(1-34)-treated cells revealed that *Vdr* and *Wnt4* genes are regulated by SIKs and CRTCs, while others are not. Although many studies have examined PTH signaling in the osteoblast/osteocyte, ours is the first to examine the global effects of these peptides on the osteoblast transcriptome. Further delineation of which signaling events are attributable to PTH(1-34), PTHrP(1-36) or ABL exclusively and which are shared among all 3 will help improve our understanding of the effects these peptides have on the osteoblast and lead to the refinement of PTH-derived treatments for osteoporosis.

## Introduction

Teriparatide (PTH(1-34)), a recombinant form of PTH, was the first osteoanabolic therapeutic to be approved by the United States Food and Drug Administration (FDA) [1,2]. Abaloparatide (ABL), an analog of parathyroid hormone-related protein (PTHrP(1-34)), became the second FDA-approved osteoanabolic for treatment of osteoporosis and has been shown to be somewhat more effective in producing osteoanabolic outcomes compared with teriparatide; ABL resulted in higher bone mineral density (BMD) in the femurs of osteoporotic, post-menopausal women compared with teriparatide [3, 4, 5]. It has also been shown that serum CTX levels, a marker for bone resorption, were lower in patients treated with ABL compared with teriparatide, suggesting it has a lesser tendency to promote deleterious effects [5]. Despite this, neither ABL nor teriparatide have been able to overcome the anabolic window preventing the long-term efficacy of these treatments for osteoporotic patients [6].

PTH, PTHrP, and ABL all bind the same G protein-coupled receptor, parathyroid hormone receptor type 1 (PTHR1) [7]. One study found that PTH (1-34), PTHrP (1-36), and ABL act through two PTHR1 conformations named R^O^ and RG. Binding to R^O^ results in prolonged signaling, which is thought to lead to comparatively more bone resorption, while binding to RG is thought to result in more osteoanabolic signaling [8]. The study determined that ABL binds with greater selectivity to RG and concluded this represents a plausible explanation for the favorable anabolic effects of ABL treatment reported on bone compared with PTH (1-34). The differences observed in this study of PTH (1-34), PTHrP (1-36), and ABL binding to PTHR1 suggest that these findings may be reflected by differential signaling events downstream of PTHR1. However, there is a paucity of data that has determined differences in signaling from PTH (1-34), PTHrP (1-36), and ABL in osteoblast lineages.

There are a variety of signaling cascades that are stimulated upon PTHR1 ligand binding. The G_Sα_/cAMP/PKA pathway accounts for most of PTHR1 signaling and this is purported to mediate the anabolic response to PTH as well as the catabolic response [9, 10, 11, 12]. Recent work from our laboratory examined if PTH (1-34), PTHrP (1-36), and ABL treatment would result in unique stimulatory effects with respect to cAMP production and known downstream effectors of this pathway, the salt-inducible kinases (SIKs) and cAMP-regulated transcriptional coactivators (CRTCs) [13]. We found that in primary murine calvarial osteoblasts, PTHrP (1-36) and ABL result in a significantly lower cAMP response compared with PTH (1-34). Downstream of this signaling cascade, time course and dose response analyses showed similar relative differences in PKA activation and the phosphorylation of cAMP response element binding protein (CREB). However, quantitative real-time PCR (qRT-PCR) of known osteoblastic genes found several genes were similarly regulated such as the Wnt inhibitor, *Sost*, while *c-Fos* and *Rankl* were differentially regulated in time and dose-dependent manners.

Since we determined that these peptides differentially exert time and dose-dependent responses in the osteoblast, particularly on the cAMP/PKA/*Fos* or *Rankl* axes, we hypothesized that these changes reflect the ability of PTH (1-34), PTHrP (1-36), and ABL to differentially modulate the osteoblast transcriptome. Many studies have examined PTH and PTHrP regulation of gene expression in stromal cells/osteoblasts/osteocytes but none to date have compared the global effects of these three peptides on the osteoblast transcriptome [14, 15, 16, 17]. In this study, RNA-sequencing was performed on primary calvarial murine osteoblasts treated with PTH (1-34), PTHrP (1-36), and ABL by repeating the conditions under which we observed differences in *Rankl* expression to compare their global effects on the osteoblast transcriptome and gene ontology [13]. Select findings were confirmed via qRT-PCR of additional cultured samples of mouse calvarial osteoblasts, treated with 1 nM of PTH (1-34), PTHrP (1-36), or ABL for 4 h prior to harvest. Lastly, analyses were performed where SIK1, SIK2, SIK3, CRTC1, CRTC2, and CRTC3 were knocked down in cells treated with 10 nM of PTH (1-34) to examine possible intertwined/separate cAMP/SIK/CRTC-dependent regulation of a number of these genes.

We hypothesized that the differences among these peptides published previously on their binding affinities, clinical outcomes, and downstream expression of anabolic/catabolic effectors would be reflected at the transcriptional level and that unique differences would be elicited by each peptide, allowing us to determine new signaling biases of interest. If these peptides lead to differential expression of osteoblastic or unknown cascades/genes then it may provide key insights for future studies to better understand the osteoanabolic and catabolic effects of PTHR1-derived treatments so that more effective therapeutics can be developed to treat osteoporosis.

## Materials and Methods

### Peptides and Chemicals

Rat parathyroid hormone (PTH 1-34) was purchased from Bachem. Parathyroid hormone-related protein (PTHrP 1-36) and abaloparatide (ABL) were synthesized by the Peptide/Protein Core Facility at the Massachusetts General Hospital (MGH). All PTH (1-34), PTHrP (1-36), and ABL peptide sequences were confirmed and analyzed for purity and degradation by NYU Grossman School of Medicine Mass Spectrometry Core Facility. All peptides were dissolved in 10 mM acetic acid. Ascorbic acid was purchased from Sigma. Collagenase A was purchased from Worthington Biochemical Corporation.

### Cell Culture

Primary mouse calvarial osteoblasts were harvested from C57Bl/6J wild-type mice aged 2-3 days postnatal. Mice were euthanized with ketamine (0.25 mg/pup) and xylazine (0.025 mg/pup). All procedures with mice were performed in accordance with an approved protocol of the Institutional Animal Care and Use Committee of New York University Grossman School of Medicine. Calvariae were digested in 1 mg/mL collagenase A at 37°C by five sequential digestions, and cells from digests 3-5 were collected and plated at a density of 6.4 x 10^3^ cells/cm^2^ in αMEM supplemented with 10% FBS, 100 units/mL penicillin, 100 ug/mL streptomycin and 0.25 μg/mL of amphotericin b. After reaching confluence, osteogenic medium (50 ug/mL ascorbic acid) was added for 5 days to allow osteoblastic differentiation. Prior to harvest, osteoblasts were treated with 1 or 10 nM of the peptides for 4 h (n=3 per group). Prior to treatment, cells were serum-starved with 0.1% FBS for 16 h.

### siRNA knockdowns

Primary mouse calvarial osteoblasts were harvested from C57Bl/6J wild-type mice aged 2-3 days postnatal and plated using the same protocol as above until they reached 70-80% confluence. The cells were given siRNAs in Lipofectamine RNAiMAX for 48 h in differentiation medium. Manufacturer’s protocol was followed such that 40 pmol of siRNA was used per well (6 well plates) with 8 µL of Lipofectamine. After 48 h of siRNA transfection, the cells were then treated with or without PTH (1-34) at 10 nM for 4 h and RNA was harvested using TRIzol™ reagent. Additional samples were collected for protein analysis after isolation with RIPA Buffer. Confirmation of siRNA knockdowns was conducted with qRT-PCR examining mRNAs, and Western blots for proteins. Relative expression of *Rpl13a, Alpl*, and *Col1a1* were examined to determine if SIK or CRTC knockdowns affected cell viability and general processes. In general, cells tolerated all knockdowns well and did not die or alter their housekeeping or cell-specific genes significantly.

### RNA-Sequencing

Total RNA was isolated from cells by using TRIzol reagent (Thermo Scientific, Pittsburgh, PA) and purified with RNeasy mini kit from Qiagen (Valencia, CA). Prior to RNA-seq, the RNA integrity was assessed with Agilent 2100 Bioanalyzer (Santa Clara, CA) and the best quality triplicate samples were chosen for the subsequent analyses. The RNA-seq libraries were constructed using the Illumina TruSeq Stranded Total RNA library prep kit with Ribozero Gold (San Diego, CA). Sequencing was carried out with an Illumina HiSeq 2500 system with paired-end 100 bp reads at the Genome Technology Center of NYU Grossman School of Medicine. The quality of raw data was checked by FastQC (v. 0.11.9), and the read counts were quantified using Salmon (v. 1.7.0) against the GRCm38/mm10 mouse transcriptome reference (UCSC) database [18]. Pairwise differential expression analysis was performed by DESeq2 R/Bioconductor package (v. 1.34.0) [19]. The data have been submitted to GEO and have the accession number GSE240235. Where gene expression was found to be significantly above ±1.0 log2FC after treatments, these genes were imputed in DAVID Bioinformatics Database NIAID/NIH for Gene Ontology (GO) analyses [20].

### qRT-PCR

Total RNA was extracted using Trizol (Sigma). Complementary DNA (cDNA) was synthesized from 1 ug of total RNA using TaqMan reverse transcription kit (Applied Biosystems) with hexamer primers following the protocol described by the manufacturer. Gene expression levels were measured using SYBR Green PCR Reagents (Applied Biosystems). The quantity of messenger RNA (mRNA) was calculated by normalizing the threshold cycle value (Ct) of specific genes to the Ct of the housekeeping genes β*-actin* and/or *Ribosomal protein L13a* (*Rpl13a*).

### Statistics

Statistical differences were analyzed either by Student’s t-test or one-way ANOVA using IBM SPSS (v24). Tukey’s tests were then performed to determine which groups in the sample differed significantly from one another. Results are expressed as mean ± S.D. and a p<0.05 was considered significant comparing treatment groups.

## Results

### Global Expression Profiles for PTH (1-34), PTHrP (1-36), and ABL

Genes were selected if they had a Log_2_ fold change (FC) > 1 and had a false discovery rate (FDR) < 0.05. RNA-Sequencing revealed that PTH (1-34) regulated 367 genes, 194 were unique; PTHrP (1-36) regulated 116 genes, 15 were unique; ABL regulated 179 genes, 20 were unique. There were 74 genes shared only among PTH(1-34) and ABL; 16 genes shared only among PTH (1-34) and PTHrP; and 83 genes shared among all three peptides. These data were analyzed and compiled into Venn diagrams (Fig.1A), heat maps comparing peptides (Fig. 1B), and volcano plots (Fig. 2). The heat maps show that there is a complete change in gene expression with PTH (1-34) and that this peptide causes greater effects than the other peptides, with PTHrP causing the least effects. The volcano plots further illustrate this and show some of the specific genes regulated and that many of the highest regulated genes are the same for the three peptides but often differ in degree of regulation.

**Figure 1:**
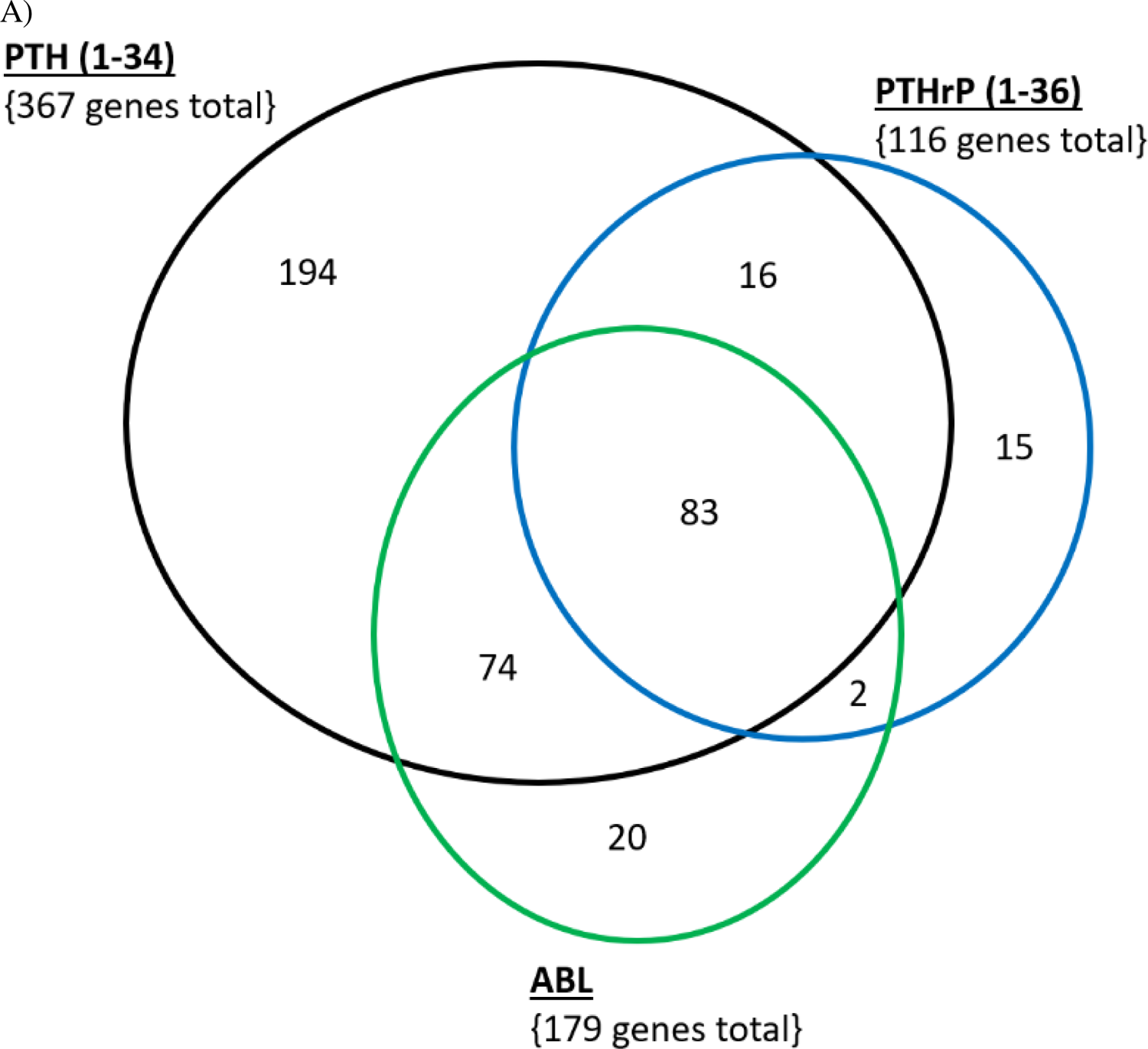
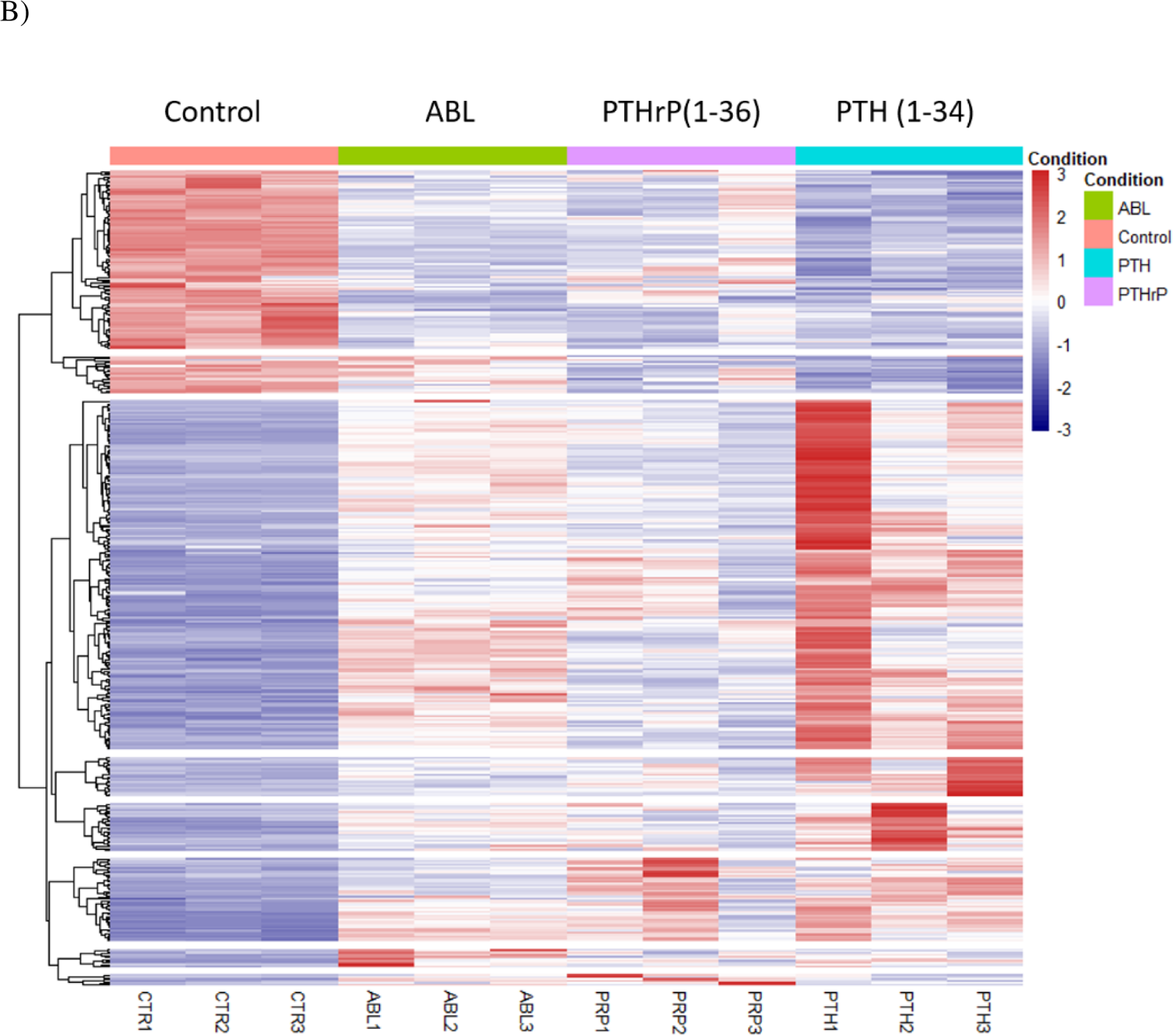
Venn Diagram and heat map of PTH (1-34), PTHrP (1-36), and ABL Treatment on the Osteoblast Transcriptome. Primary mouse calvarial osteoblasts were treated with 1 nM of these peptides for 4 h and gene enrichment analysis of RNA-Seq data was performed. Gene lists were selected with Log_2_ Fold Change ≥ 1; False Discovery Rate < 0.05 compared with the vehicle controls. A) PTH (1-34) regulated 367 genes, 194 were unique; PTHrP (1-36) regulated 116 genes, 15 were unique; ABL regulated 179 genes, 20 were unique. There were 74 genes shared only among PTH(1-34) and ABL; 16 genes shared only among PTH (1-34) and PTHrP; and 83 genes shared among all three peptides. B) Red represents transcript upregulation; blue represents transcript downregulation. Analysis is relative to control samples. Genes were selected if they had a Log_2_ fold change (FC) > 1 and had a false discovery rate (FDR) < 0.05.

**Figure 2:**
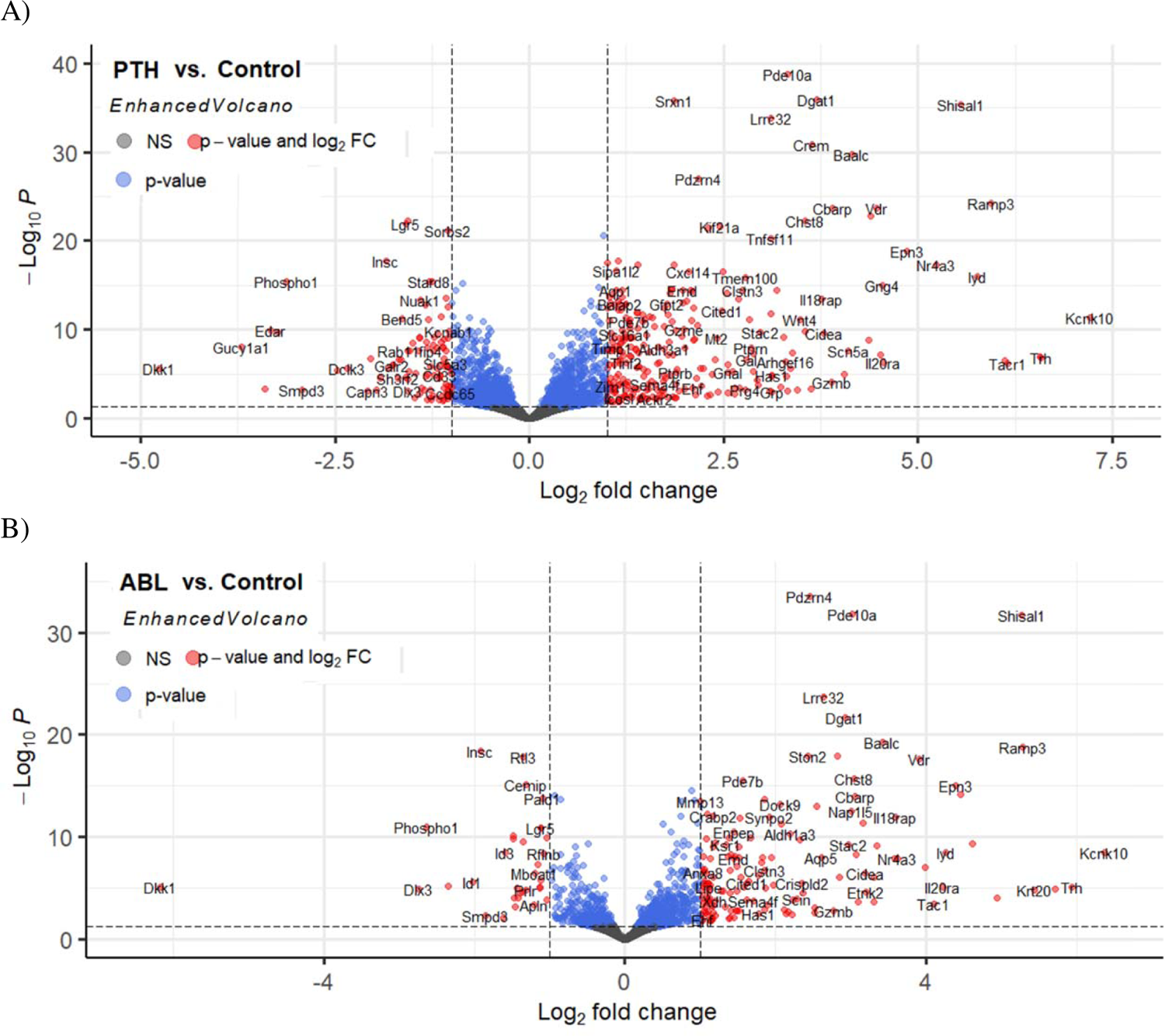
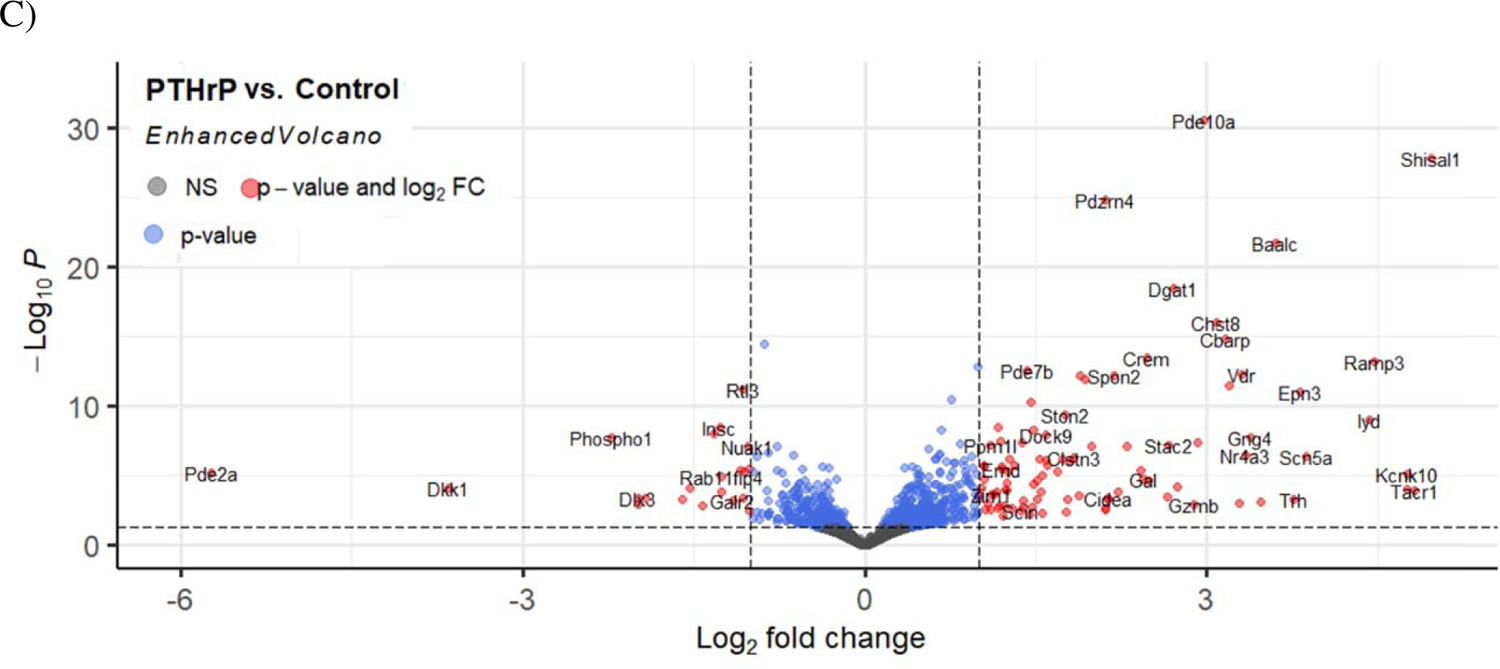
Volcano Plots of PTH (1-34), PTHrP (1-36), and ABL-regulation of the Osteoblast Transcriptome. Primary mouse calvarial osteoblasts were treated with 1 nM of these peptides for 4 h and gene enrichment analysis of RNA-Seq data was performed. Genes were selected if they had a Log_2_ fold change (FC) > 1 and had a false discovery rate (FDR) < 0.05. (A) Shows PTH (1-34) vs. Control, (B) ABL vs. Control, (C) PTHrP (1-36) vs. Control. Genes which are red on the graph are significantly regulated and have a Log_2_ fold change (FC) > 1; genes represented by blue were significantly regulated but did not pass the FC threshold, and genes represented by grey had no significant changes compared to control with either selection.

### Pathway Analyses of PTH (1-34), PTHrP (1-36), and ABL Treatment on the Osteoblast Transcriptome

Analyses of RNA-seq data were performed using gene ontology (GO) to determine pathway-specific trends from differentially expressed genes according to each peptide treatment (Fig. 3). These were analyzed via R to identify the gene ontology biological processes, molecular functions and cellular components affected by the three peptides. In this figure the size of the bubbles corresponds to the number of genes each peptide regulated per category and the color represents the level of significance. With respect to biological processes, PTH (1-34) and ABL have similarly high levels of significance for all categories except for “morphogenesis of branching structures”, where ABL does not significantly regulate this process, while PTHrP (1-36) and PTH (1-34) showed more common regulation of genes of this process. PTH (1-34) and ABL show an almost identical pattern otherwise, and PTHrP (1-36) regulates these processes differently when compared with either of these peptides. Some processes were regulated by all three, some by just two. Notably, PTH (1-34) and ABL similarly regulated genes of “bone mineralization”, “biomineral tissue development” and “actin filament bundle organization”, while PTHrP (1-36) did not regulate these processes significantly. Many molecular functions were highly regulated by PTH (1-34), e.g., receptor ligand activity, transcription factor activity, phospholipid binding, cytokine and receptor activity. Several were similarly regulated by all three peptides; phosphoric ester hydrolase activity, G-protein receptor activity and nuclear receptor activity. Likewise, PTH (1-34) generally gave the greatest and most significant regulation of all the cell components regulated, such as receptor complex, membrane rafts and collagen-containing extracellular matrix. PTHrP yielded the least effects for both molecular functions and cell components.

**Figure 3:**
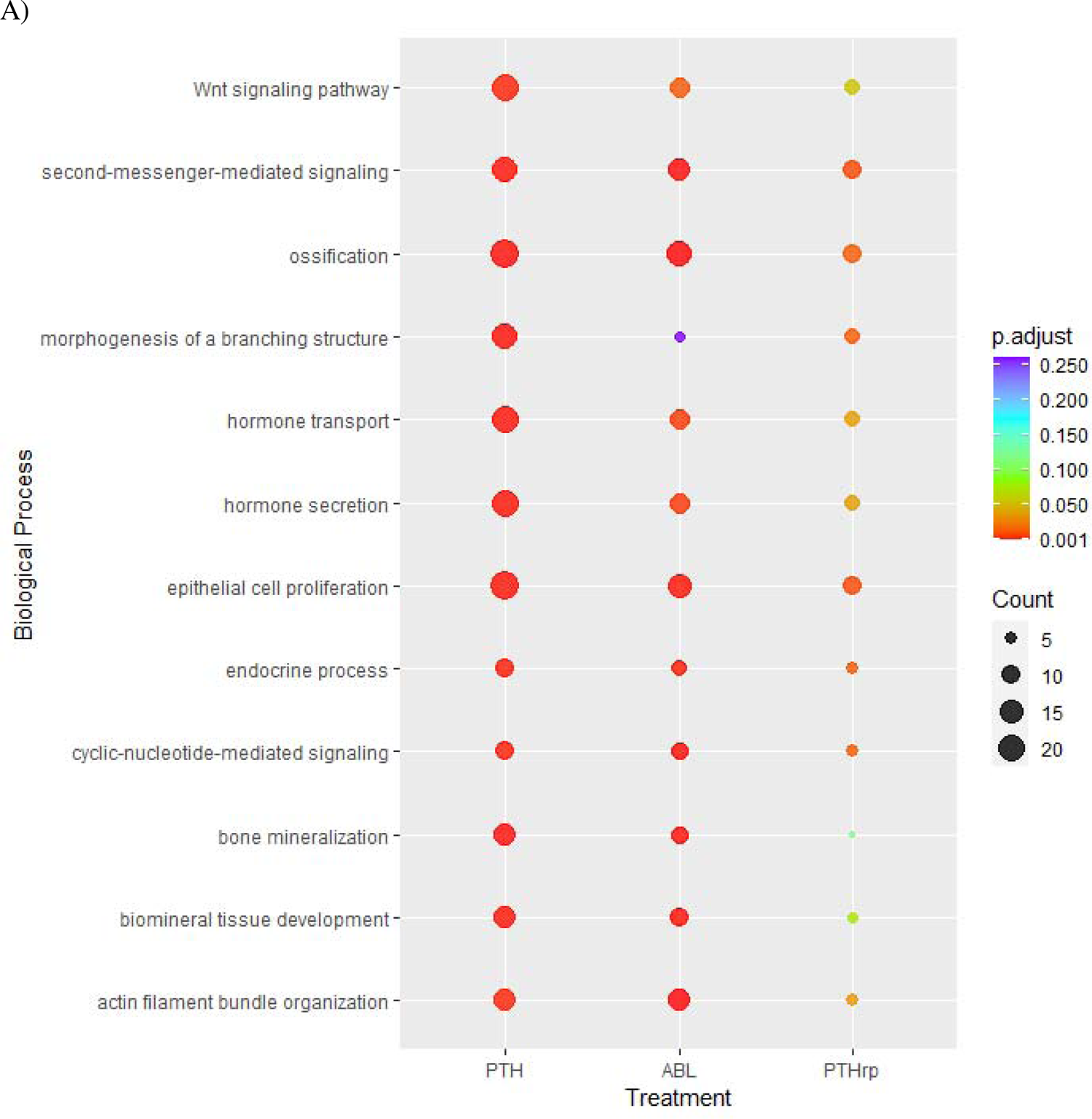
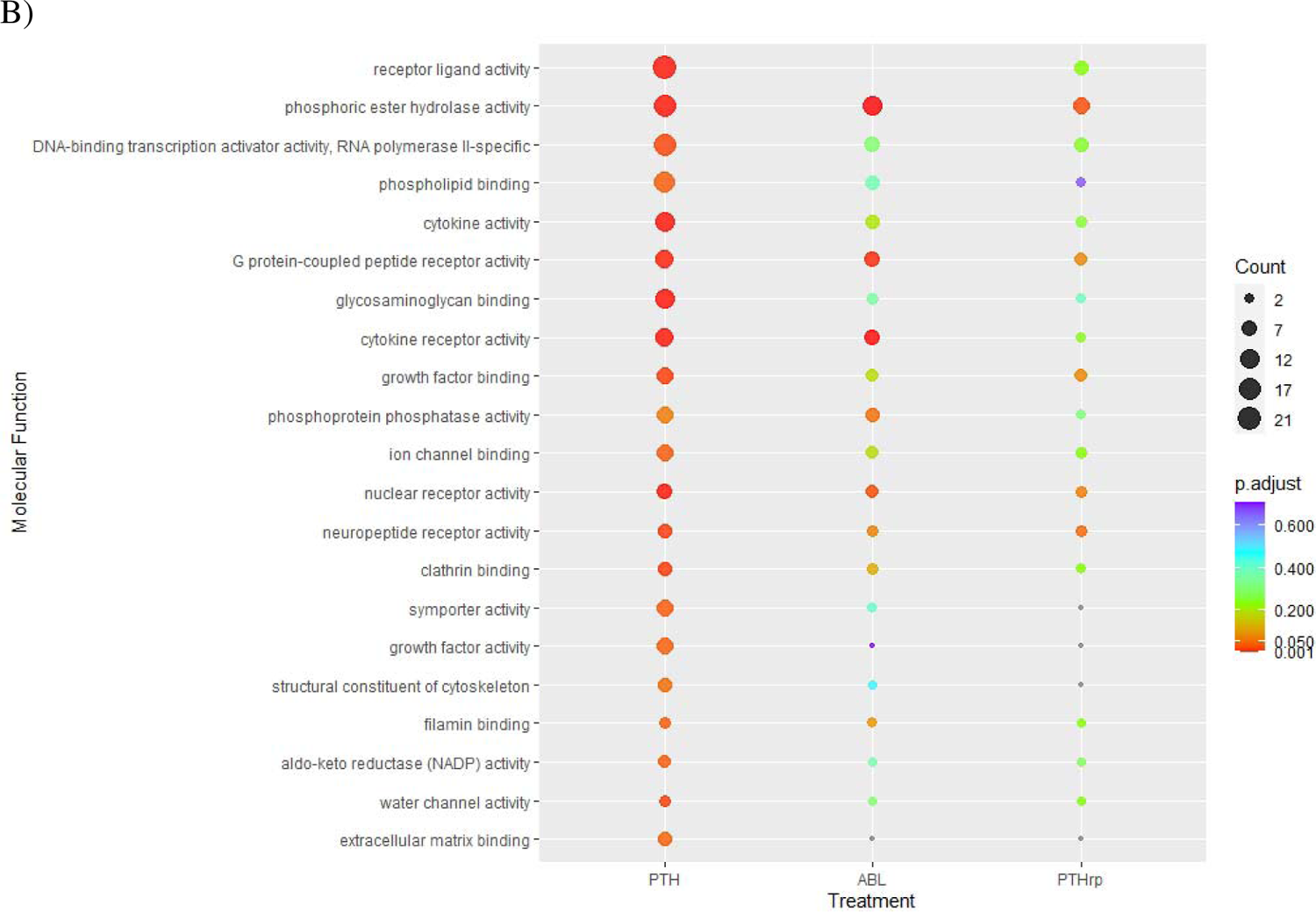
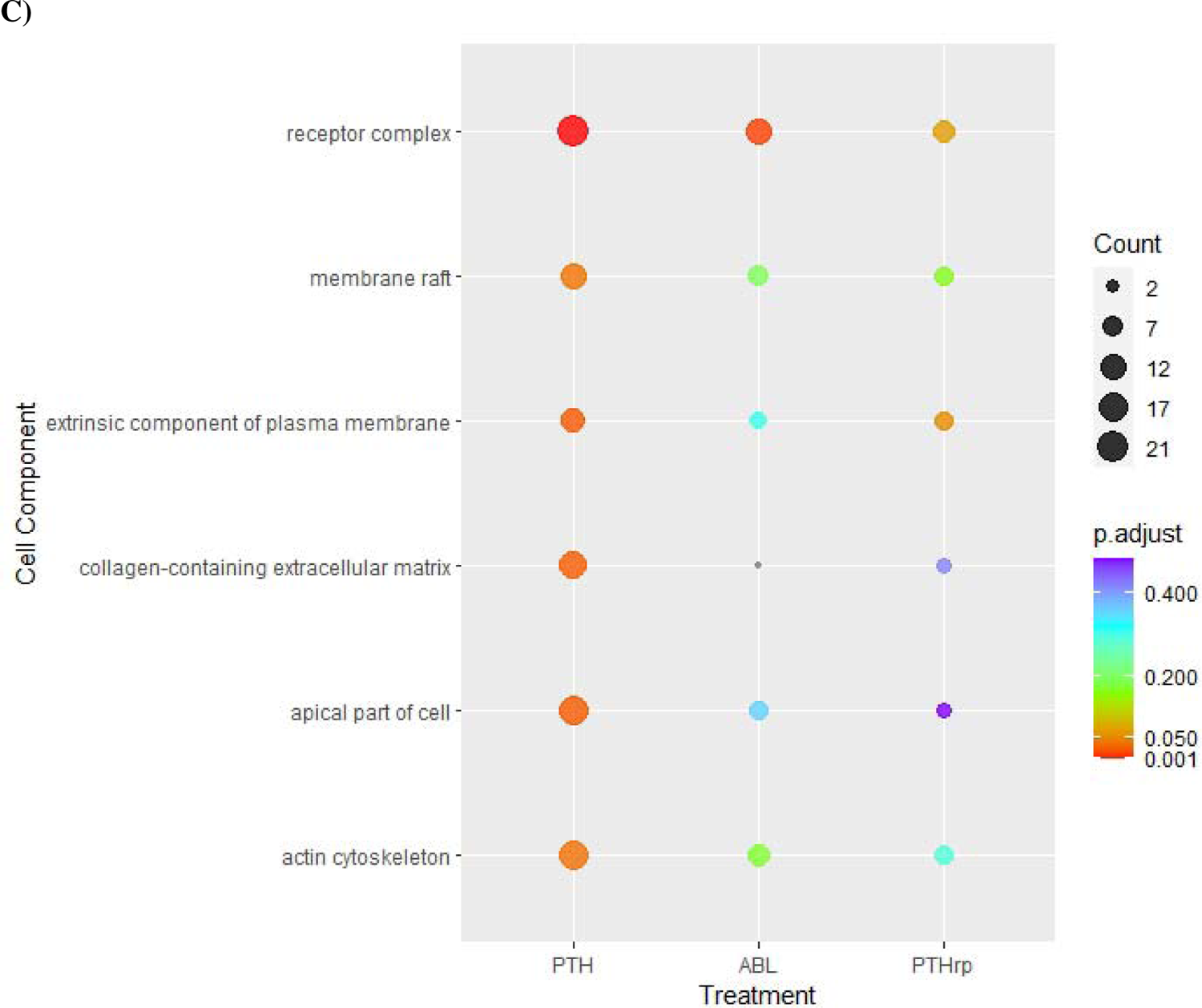
Pathway Specific Comparison of PTH (1-34), ABL, and PTHrP (1-36) Treatment on Osteoblast Transcriptome Pathways. Primary mouse calvarial osteoblasts were treated with 1 nM of these peptides for 4 h and gene enrichment analysis of RNA-Seq data was performed. If gene expression was found to be above ±1.0 log_2_FC genes were imputed in DAVID Bioinformatics Database NIAID/NIH for Gene Ontology (GO) analysis; data for Control vs. peptide were graphed via R.

### Examination of genes of interest with RNA-seq data and qRT-PCR verification

Based on these data we selected several genes which were highly up or down-regulated to examine more closely and to verify the RNA-seq data with qRT-PCR in separate calvarial osteoblast preparations. The genes included 1) *Cbp/P300 interacting transactivator with Glu/Asp rich carboxy-terminal domain 1* (*Cited1*), 2) *Vitamin D receptor* (*Vdr*), 3) *Phosphodiesterase 10A* (*Pde10a*), 4) *Wnt Family Member 11* (*Wnt11*), 5) *Secreted frizzled related protein 4* (*Sfrp4*), 6) *cAMP responsive element modulator* (*Crem*), 7) *Ephrin type-A receptor 3* (*Epha3*), 8) *Histone deacetylase 4 (Hdac4*), 9) *Protein phosphatase 2 regulatory subunit B alpha* (*Pp2r2a*), 10) *Dishevelled segment polarity protein 3* (*Dvl3*), 11) *Wnt4*, 12) *Wnt7b*, 13) *Dickkopf1, Dkk1*, 14) *Frizzled class receptor 5* (*Fzd5*), 15) *Transcription factor 7* (*T-Cell specific, HMG-box*) (*Tcf7*), and 16) *Potassium two pore domain channel subfamily K member 10* (*Kcnk10*). RNA-seq Log_2_ fold changes/p-values are shown in Table 1.

**Table 1:** Regulation of genes of interest by RNAseq after PTH (1-34), ABL, and PTHrP (1-36) treatment in differentiating calvarial osteoblasts. Primary mouse calvarial osteoblasts were treated with 1 nM of these peptides for 4 h and gene enrichment analysis of RNA-Seq data was performed. Gene lists were selected with Log_2_ Fold Change ≥ 1; False Discovery Rate < 0.05 compared with the vehicle controls. (+) log2 FC represents upregulation compared to control; (-) log2 FC represents downregulation compared to control.

| Gene ID | Gene Name | PTH (1-34) |  | ABL |  | PTHrP (1-36) |  |
| --- | --- | --- | --- | --- | --- | --- | --- |
|  |  | Log2 FC | p-value | Log2 FC | p-value | Log2 FC | p-value |
| **Cited1** | Cbp/P300 Interacting Transactivator 1 | 2.48 | 3.56E-15 | 1.61 | 3.22E-08 | 1.56 | 7.13E-08 |
| **Vdr** | Vitamin d receptor | 4.47 | 2.31E-24 | 3.90 | 1.83E-21 | 3.31 | 4.77E-16 |
| **Pde10a** | Phosphodiesterase 10A | 3.34 | 1.02E-43 | 3.02 | 2.21E-36 | 2.96 | 2.10E-35 |
| **Wnt11** | Wnt family member 11 | 1.52 | 1.43E-07 | 1.15 | 1.27E-05 | 0.13 | ns |
| **Sfrp4** | Secreted Frizzled Related Protein 4 | 1.141 | 3.00E-07 | 1.054 | 3.77E-06 | 0.355 | 4.09E-02 |
| **Crem** | cAMP responsive element modulator | 2.30 | 0.0001 | 2.82 | 8.53E-22 | 2.47 | 2.21E-17 |
| **Epha3** | ephrin type-A receptor 3 | 4.86 | 3.11E-06 | 4.35 | 5.14E-11 | 4.71 | 4.65E-05 |
| **Hdac4** | Histone deacetylase 4 | 1.33 | 4.56E-14 | 1.20 | 1.33E-12 | 1.16 | 5.44E-12 |
| **Pp2r2a** | Protein Phosphatase 2 Regulatory Subunit B alpha | 0.205 | 2.88E-01 | 0.013 | 9.43E-01 | 0.393 | 5.23E-02 |
| **Dvl3** | Dishevelled Segment Polarity Protein 3 | 0.092 | 6.42E-01 | 0.035 | 8.17E-01 | 0.756 | 1.14E-02 |
| **Wnt4** | Wnt family member 4 | 3.49 | 4.68E-14 | 3.08 | 2.67E-11 | 1.18 | 3.17E-06 |
| **Wnt7b** | Wnt family member 7b | 0.274 | 1.73E-01 | 0.397 | 6.22E-02 | 0.106 | 4.09E-01 |
| **Dkk1** | Dickkopf WNT signaling pathway inhibitor 1 | -4.77 | 2.42E-11 | -6.17 | 7.83E-08 | -3.65 | 8.04E-07 |
| **Fzd5** | Frizzled Class Receptor 5 | -1.279 | 1.32E-06 | -0.805 | 1.83E-03 | -0.154 | 2.90E-01 |
| **Tcf7** | Transcription Factor 7 (T-Cell Specific, HMG-Box) | -0.572 | 3.42E-02 | -0.128 | 3.44E-01 | -0.208 | 1.53E-01 |
| **\*\*\*\*\***<br>**Kcnk 10** | Potassium two pore domain channel subfamily K member 10 | 7.21 | 4.30E-12 | 2.83 | 8.53E-22 | 4.76 | 5.49E-08 |

qRT-PCR data confirmed *Vdr, Cited1, Pde10a*, and *Wnt11* mRNAs followed the reported pattern of expression profiling of *Rankl* in response to these peptides: PTHrP (1-36) and ABL elicit a moderate increase in *Rankl* compared with control with PTHrP (1-36) being in most cases significantly lower than ABL and PTH (1-34) eliciting the greatest increase in expression of these genes compared with control (Fig 4). *Sfrp4* closely mimicked this expression pattern but with one minor difference in just the qPCR data: PTHrP (1-36) resulted in higher mRNA expression than ABL but both were still lower than PTH (1-34) treatment.

**Figure 4:**
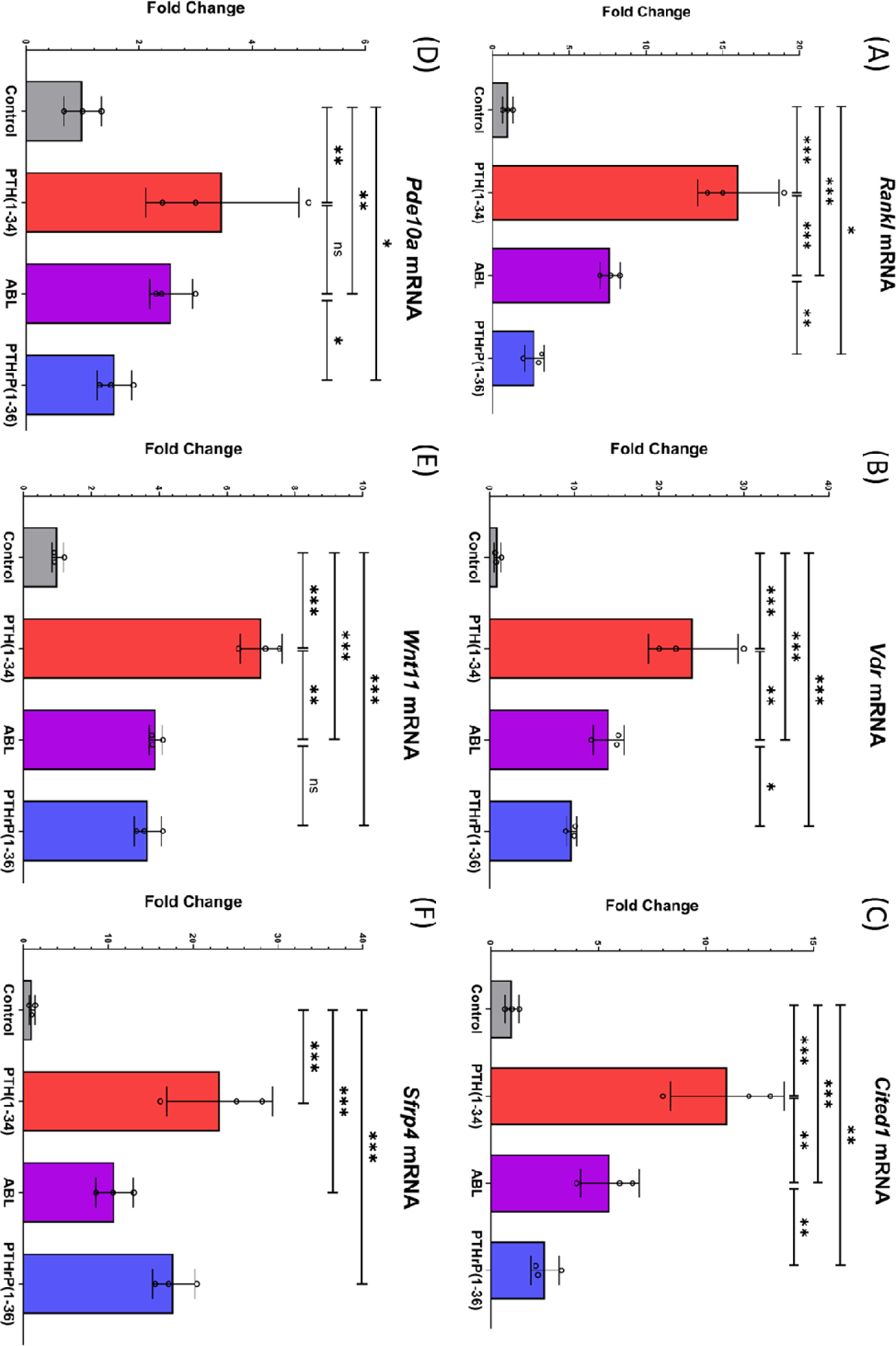
PTH (1-34), PTHrP (1-36), and ABL Modulate Expression of *Vdr, Cited1, Pde10a, Wnt11,* and *Sfrp4* mRNA with a Similar Pattern to *Rankl*. Primary differentiating calvarial osteoblasts were treated with 1 nM of PTH (1-34), PTHrP (1-36), or ABL for 4 h prior to harvest, followed by qRT-PCR for *Vdr, Cited1, Pde10a, Wnt11, Sfrp4* and *Rankl* mRNAs. All data are expressed relative to the housekeeping gene *Rpl13a* and represent mean ± S.D. of n=3 independent experiments. * p <0.05, ** p<0.01, *** p< 0.001, ns, not significantly different.

qRT-PCR data showed that *Crem, Epha3, Hdac4, Pp2r2a,* and *Dvl3* mRNAs were similarly regulated by all three peptides even if not all were significantly so (Fig 5), confirming the RNA-seq data (Table 1).

**Figure 5:**
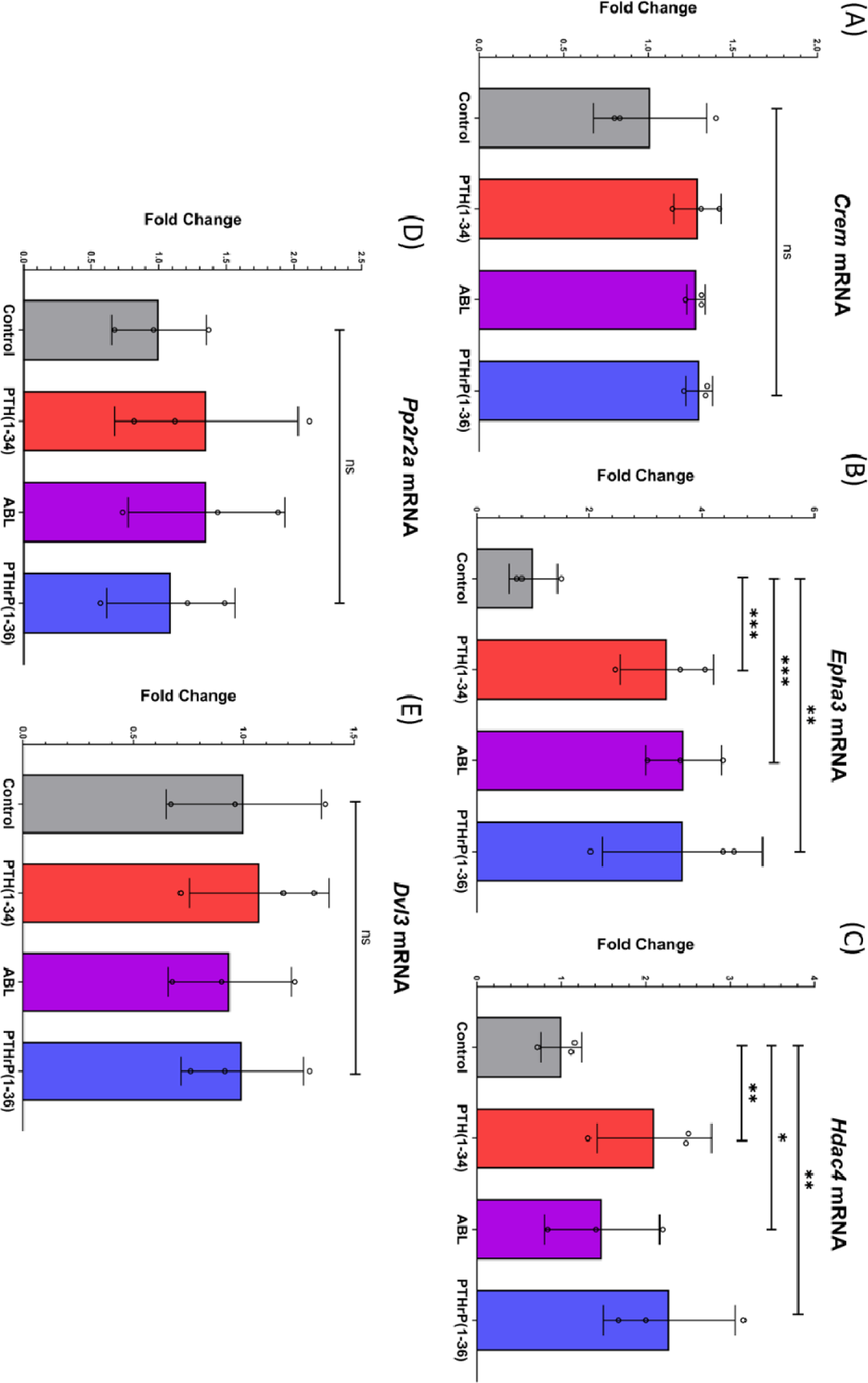
PTH (1-34), PTHrP (1-36), and ABL have the Same Effect on *Crem, Epha3, Hdac4, Pp2r2a,* and *Dvl3* mRNA Expression. Primary differentiating calvarial osteoblasts were treated with 1 nM of PTH (1-34), PTHrP (1-36), and ABL for 4 h prior to harvest, followed by qRT-PCR for *Crem, Epha3, Hdac4, Pp2r2a,* and *Dvl3* mRNAs. All data are expressed relative to the housekeeping gene β*-actin* and represent mean ± S.D. of n=3 independent experiments. * p <0.05, ** p<0.01, *** p< 0.001, ns, not significantly different.

Further qPCR analyses show additional mRNA response patterns that mimic their respective RNA-seq data bar one, *Kcnk10* (Table 1). *Wnt4* and *Wnt7b* are both upregulated by all three peptides compared to control and it is very noticeable that PTHrP yields a significantly greater stimulation of *Wnt4* than the other peptides. *Dkk1, Fzd5*, and *Tcf7* are all downregulated by all three peptides (Fig 6). These qPCR data analyses show all three peptides downregulated *Kcnk10* which differs from the RNA-seq data which shows upregulation for all three peptides, PTH (1-34) having the greatest impact of them all.

**Figure 6:**
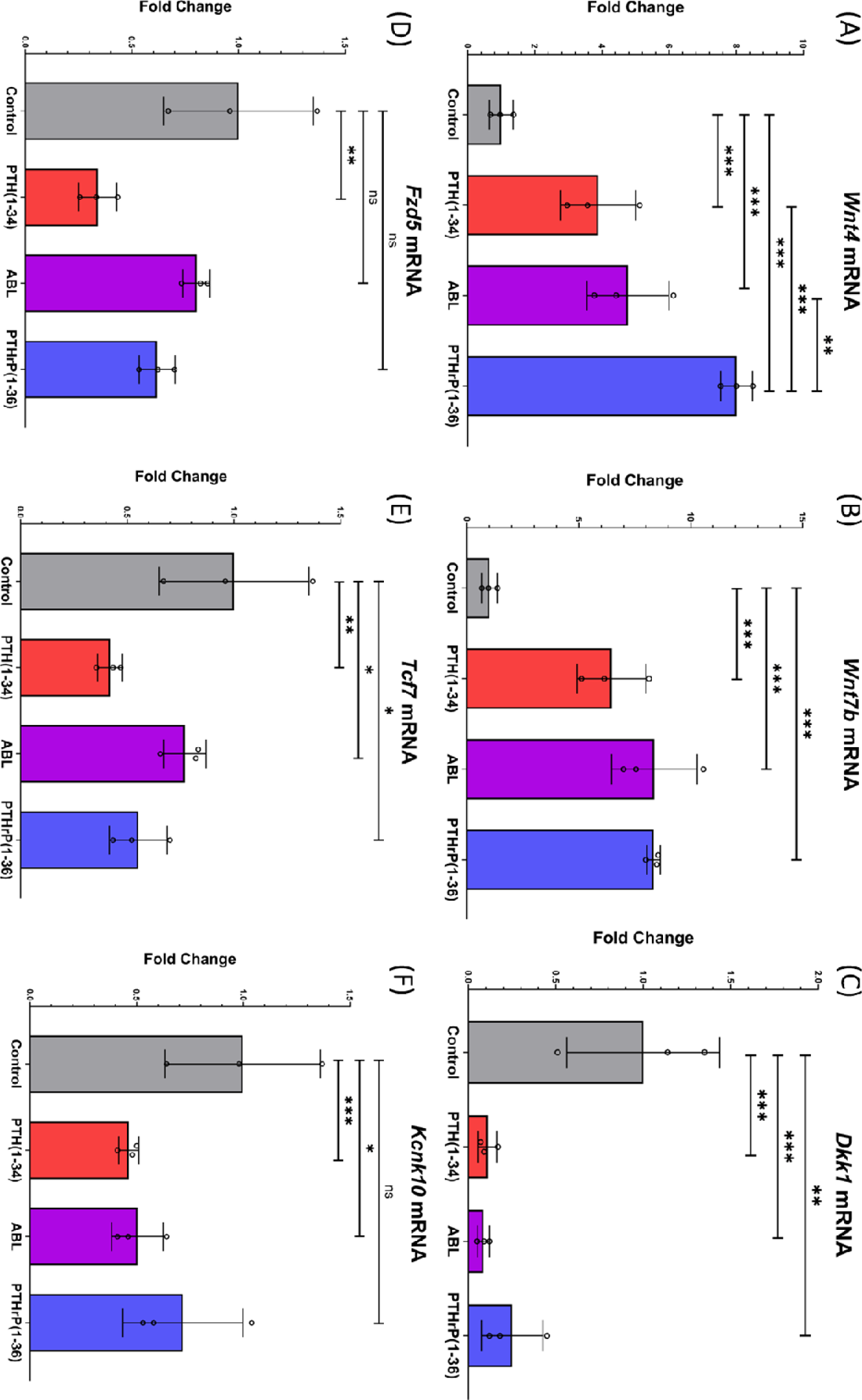
PTH (1-34), PTHrP (1-36), and ABL Regulation of *Wnt4, Wnt7b, Dkk1, Fzd5, Tcf7*, and *Kcnk10* mRNAs. Primary differentiating calvarial osteoblasts were treated with 1 nM peptides for 4 h, followed by qRT-PCR for *Wnt4, Wnt7b, Dkk1, Fzd5, Tcf7,* and *Kcnk10*. All data are expressed relative to the housekeeping gene *Rpl13a* and represent the mean ± S.D. of n=3 independent experiments. * p <0.05, ** p<0.01, *** p< 0.001, ns, not significantly different.

### Examination of genes of interest with siRNA knockdowns of SIK1, SIK2, SIK3, CRTC1, CRTC2, and CRTC3

Since several of the genes followed the same differential regulation by these peptides as *Rankl*, we investigated if the genes were regulated through the SIK/CRTC pathway, as *Rankl* is [13]. We and others have shown that PTH inhibits SIK2 and 3 resulting in dephosphorylation of CRTC2 and 3, the translocation of the latter into the nucleus and a resultant increase in *Rankl* transcription [13, 21].

*Cited1, Vdr, Wnt11, Wnt4, Wnt7b, Sfrp4, Epha3,* and *Sost* mRNA abundance were examined after SIK1/2/3 & CRTC1/2/3 were knocked down in osteoblastic cells that were then treated with PTH (1-34) (Fig 7). Knockdowns did not significantly change any of the above mRNAs under basal conditions but substantial individual/combined regulation was found in these samples when compared to scrambled controls and PTH (1-34) treatment. Analysis of the effects of SIK1/2/3 or CRTC1/2/3 knockdowns revealed complex relationships; data show that some of these genes are regulated by both SIKs and CRTCs (*Vdr, Wnt4*) as *Rankl* is, while others (*Sfrp4*) are regulated by SIKs but not CRTCs, in the same way as *Sost* is and some require SIK expression [21].

**Figure 7:**
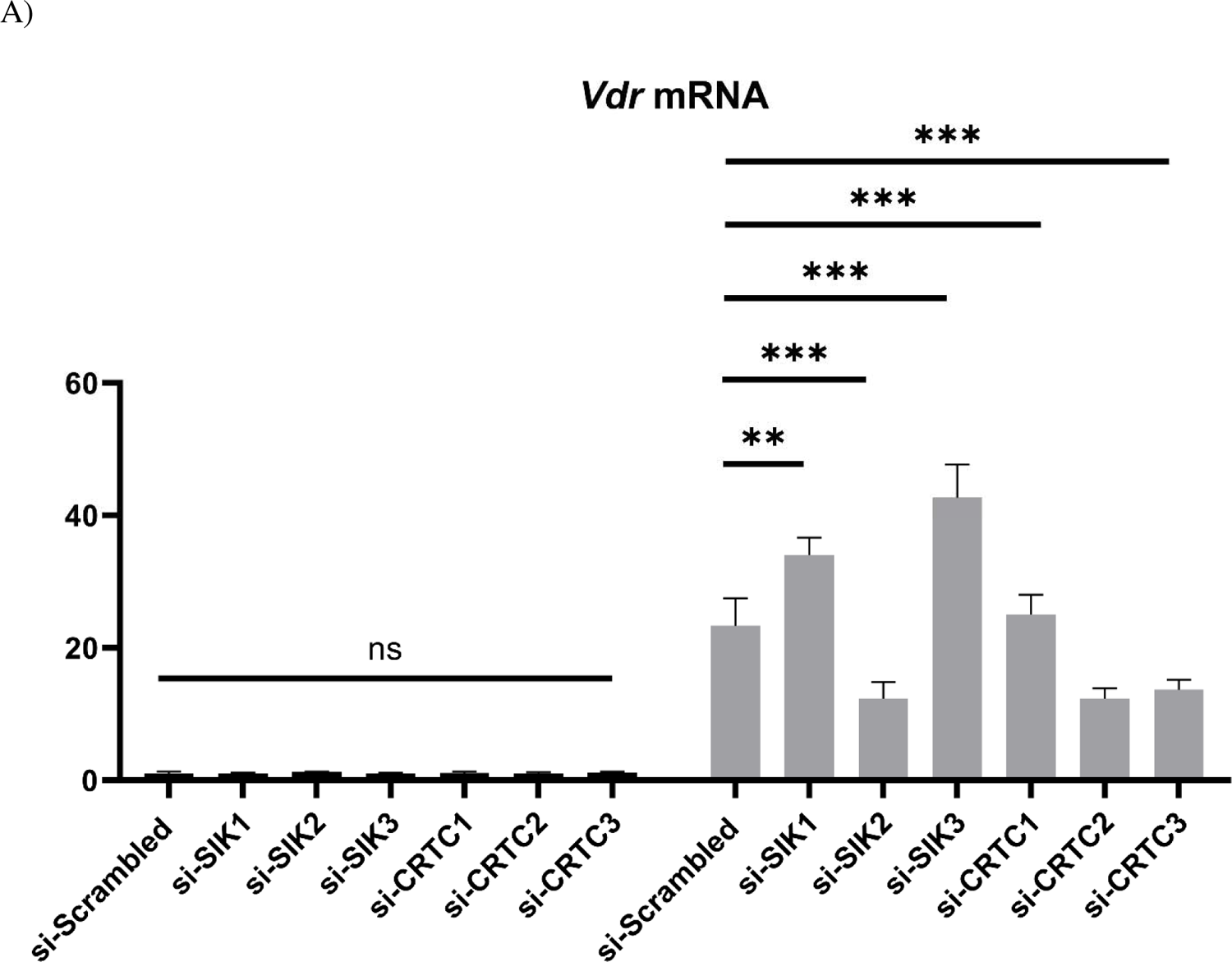
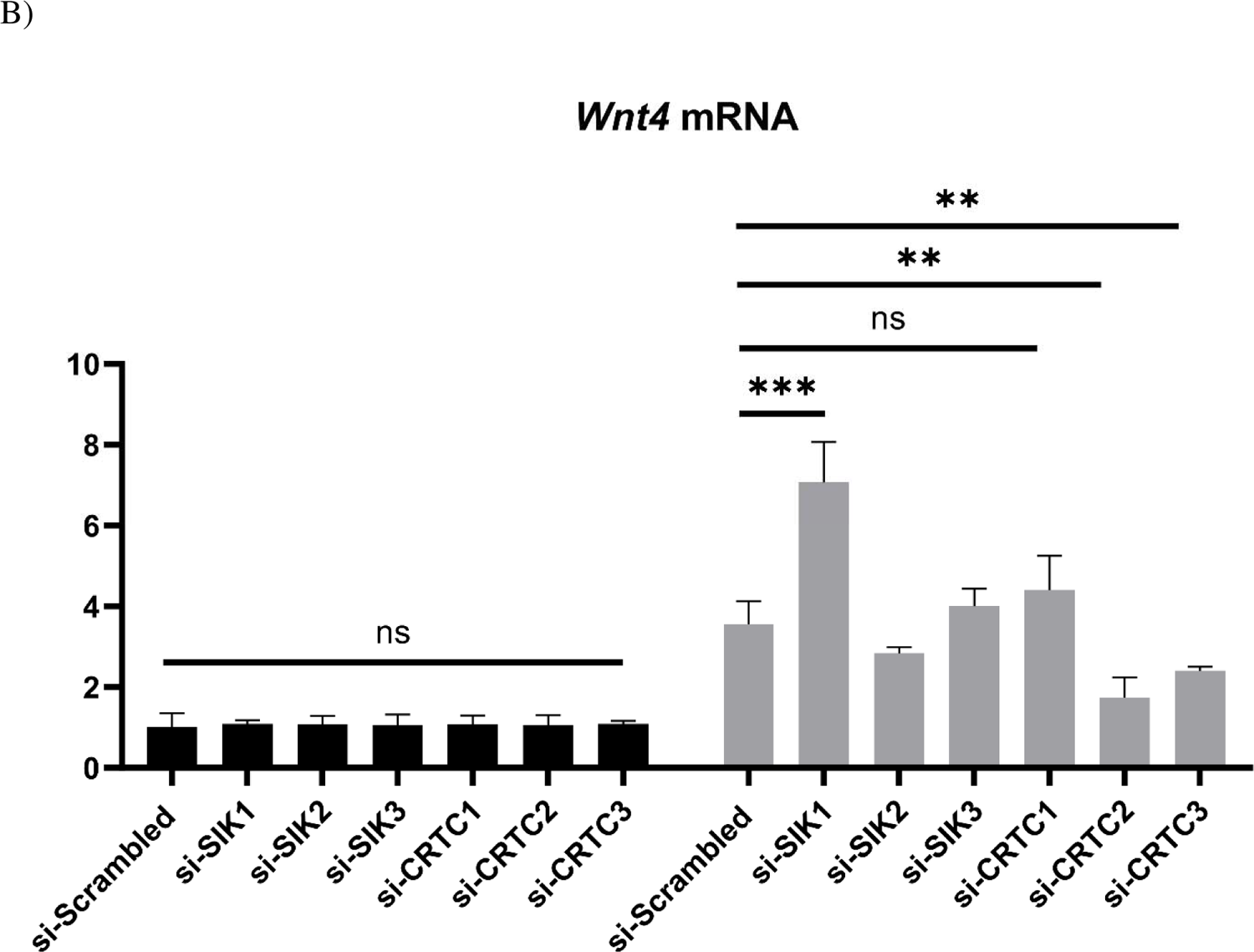
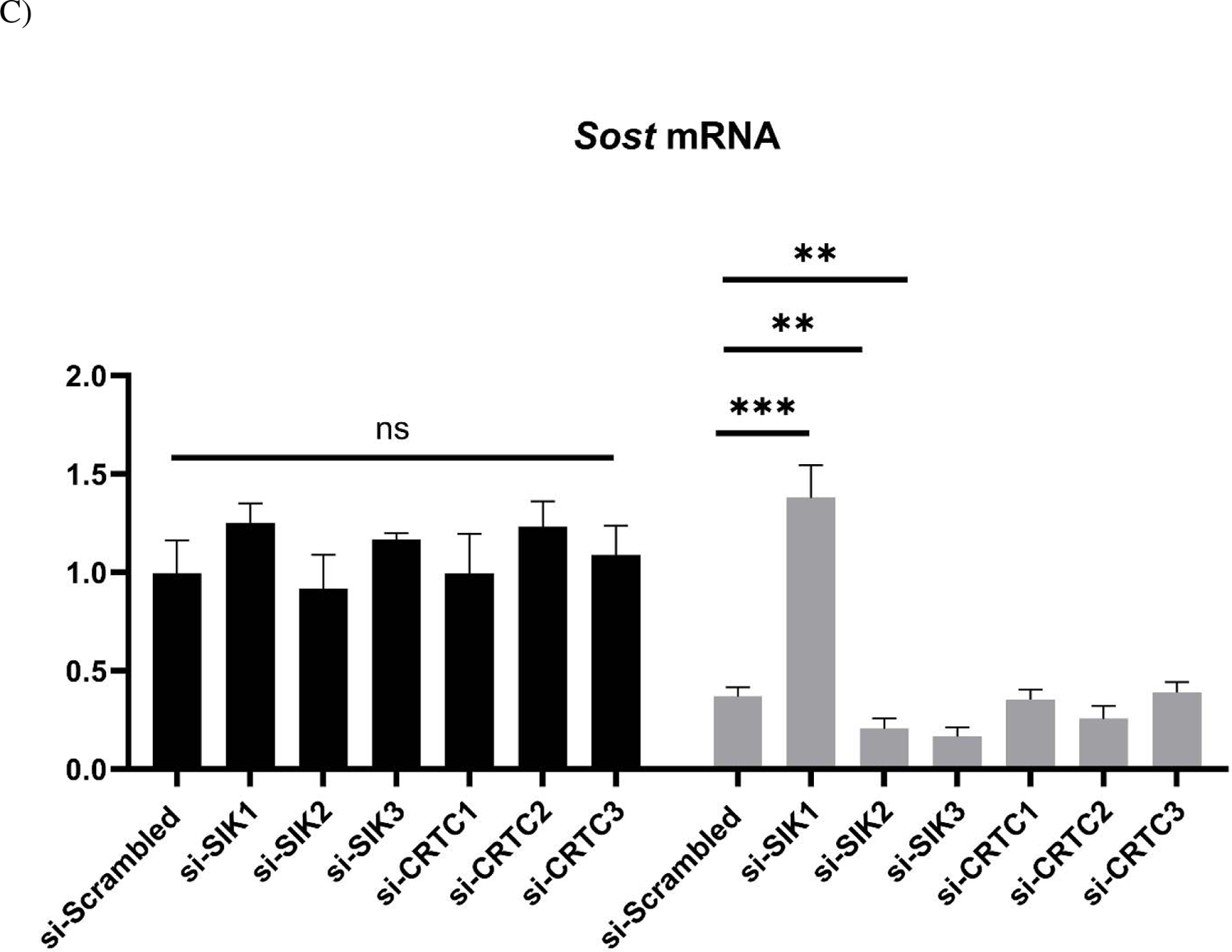
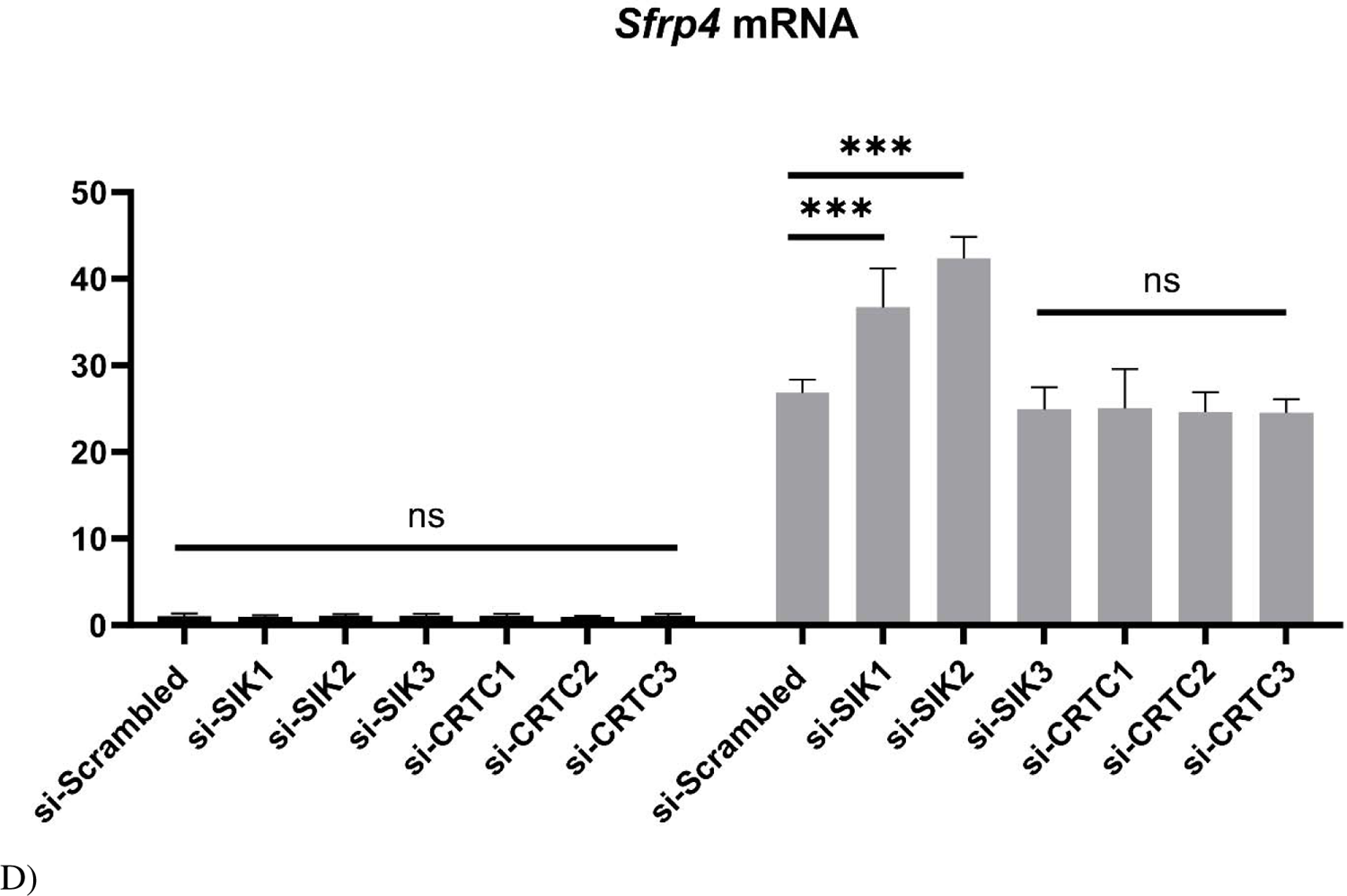
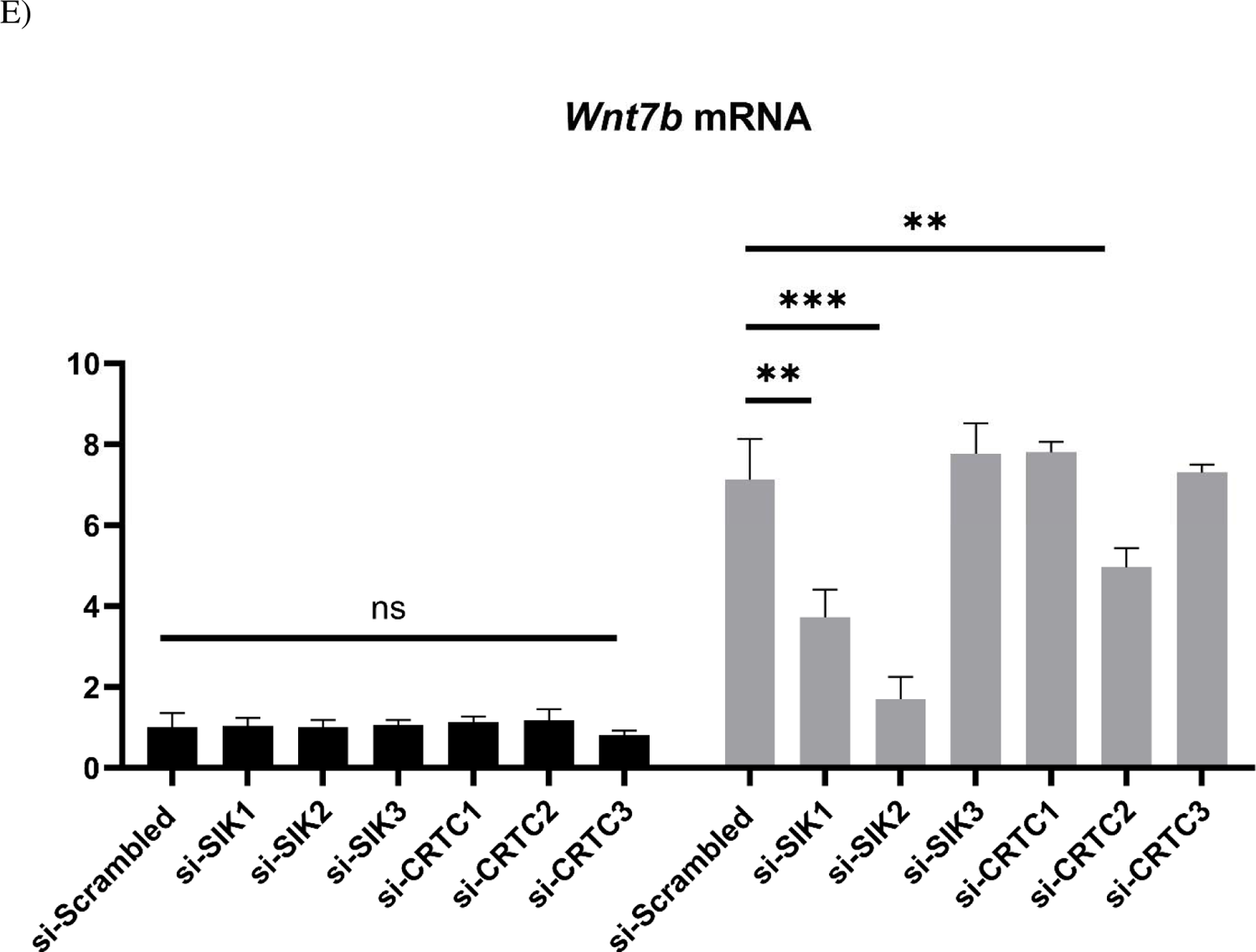
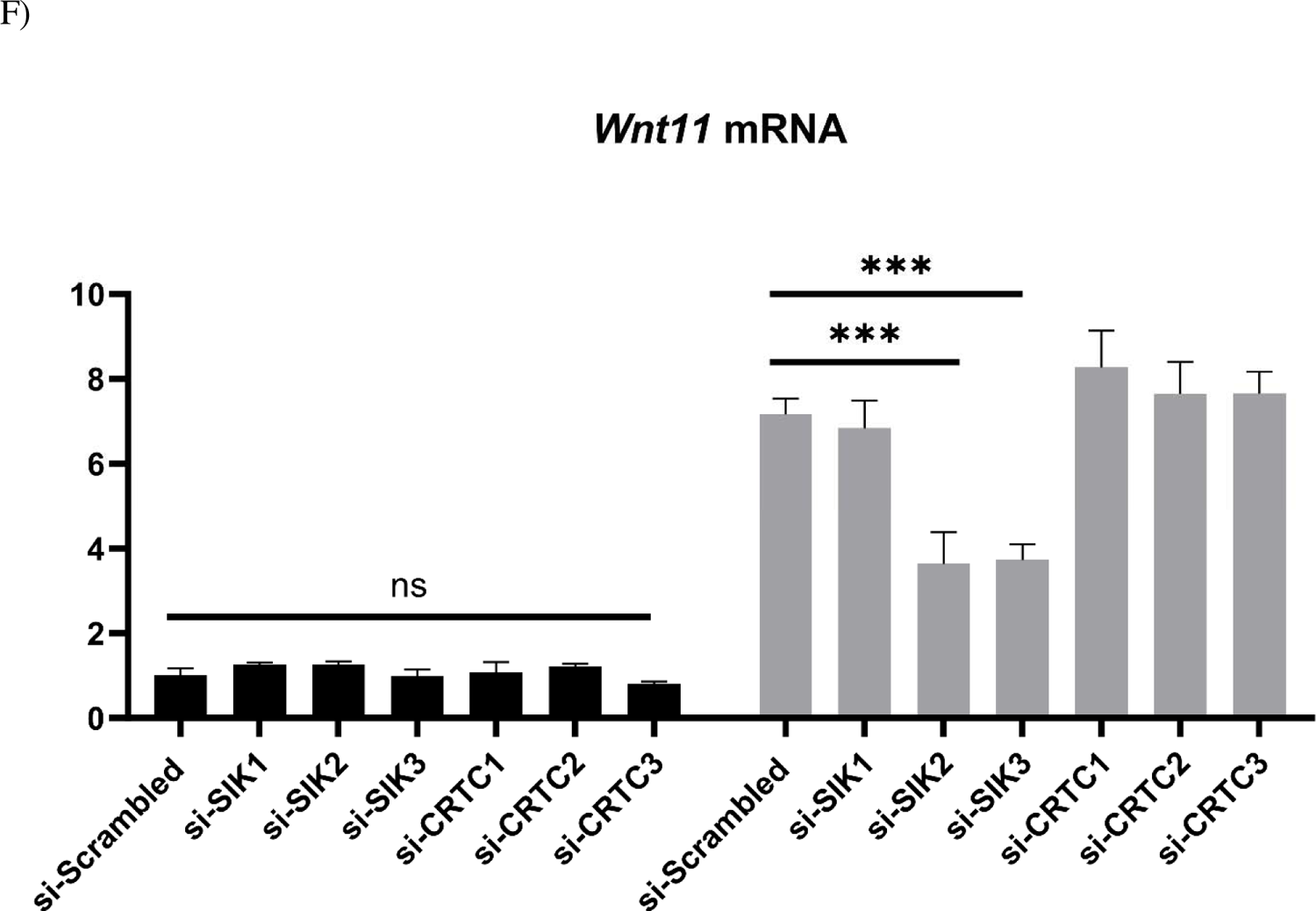
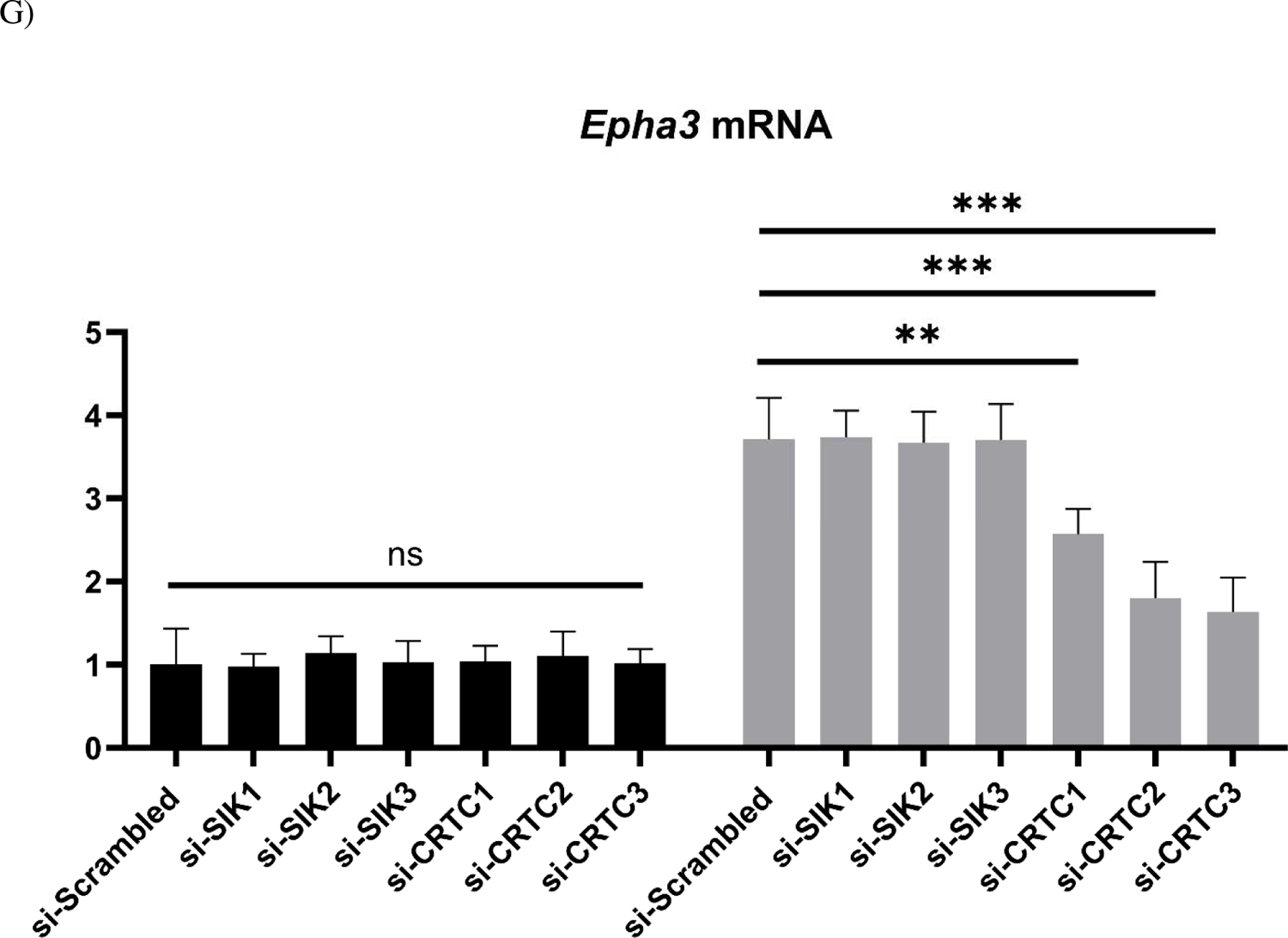
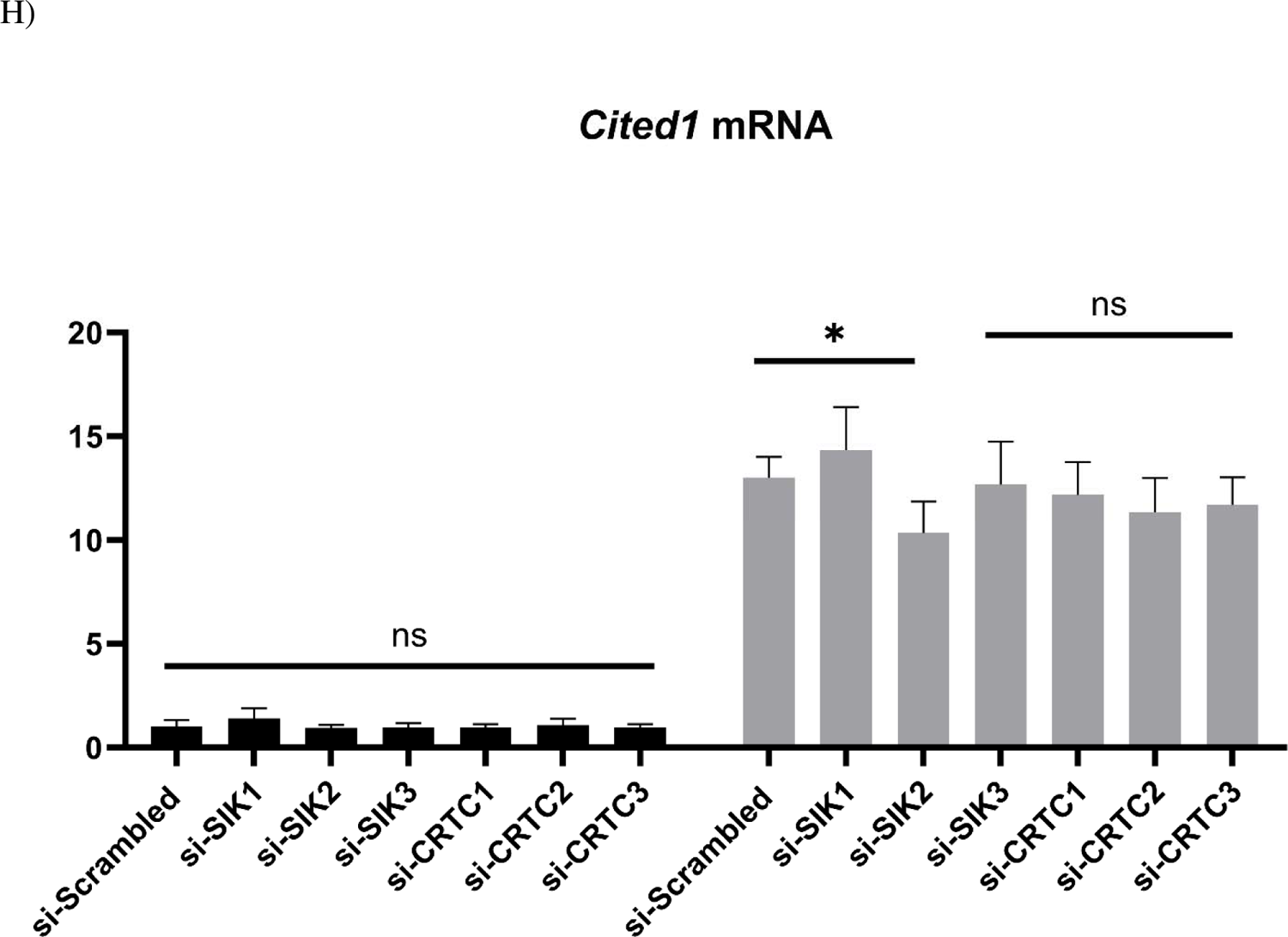
Role of SIK1/2/3 & CRTC1/2/3 in PTH (1-34) regulation of *Vdr, Wnt4, Sost, Sfrp4, Wnt7b, Wnt11, Epha3,* and *Cited1 mRNA* in mouse calvarial osteoblasts. Primary mouse calvarial osteoblasts were cultured to 70-80% confluence. The cells were given siRNAs for si-*Sik1*, si-*Sik2*, si-*Sik3*, si-*Crtc1*, si-*Crtc2*, & si-*Crtc3* (40 pmol) in lipofectamine RNAiMAX for 48 h in osteoblast differentiating medium. The cells were then treated with or without PTH (1-34) at 10 nM for 4 h; RNA was harvested to determine mRNA in basal and PTH-treated samples compared to si-scrambled controls. All data are expressed relative to the housekeeping gene *Rpl13a* and represent mean ± S.D. * p <0.05, ** p<0.01, *** p< 0.001, ns, not statistically different.

*Vdr* mRNA significantly increased with PTH treatment and SIK1 and SIK3 knockdowns when compared with scrambled controls; *Vdr* mRNA levels significantly decreased with knockdown of SIK2 and CRTC2/3 compared with scrambled controls. Knockdown of CRTC1 had no significant effect. Similarly, *Wnt4* mRNA significantly increased with PTH treatment and SIK1 knockdown when compared with scrambled controls; *Wnt4* mRNA significantly decreased with CRTC2/3 knockdown compared with scrambled controls. SIK2/3 and CRTC1 knockdowns had no statistically significant effects.

*Sost* mRNA significantly increased with SIK1 knockdown compared with scrambled controls; *Sost* mRNA significantly decreased with SIK2/3 knockdowns compared with scrambled controls. Knockdowns of CRTC1/2/3 had no statistically significant effects. *Sfrp4* mRNA significantly increased in SIK1/2 knockdowns when compared with scrambled controls; SIK3 and CRTC1/2/3 had no statistically significant effects.

*Wnt7b* mRNA significantly decreased with SIK1/2 and CRTC2 knockdowns compared with scrambled controls. SIK3 and CRTC1/3 knockdowns had no statistically significant effects. *Wnt11* mRNA significantly decreased with SIK2 and SIK3 knockdown when compared with scrambled controls; SIK1 and CRTC1/2/3 knockdowns had no statistically significant effects.

*Epha3* mRNA significantly decreased with CRTC2/3 knockdown compared with scrambled controls. SIK1/2/3 and CRTC1 knockdown had no statistically significant effects.

*Cited1* mRNA was barely affected by any of the SIK or CRTC knockdowns, suggesting it is not regulated by PTHR1 through this pathway.

## Discussion

In this study we show that PTH (1-34), PTHrP (1-36), and ABL treatment of calvarial osteoblasts *in vitro* result in differing effects on the osteoblast transcriptome using RNA-Seq, GO and additional qRT-PCR of a number of genes of interest. We further investigated the mechanism of regulation of some of these genes using SIK1/2/3 and CRTC1/2/3 knockdowns in cells treated with PTH (1-34) and found several commonly-regulated by SIK or CRTC-dependent pathways while others showed complex regulation. Previous research from our laboratory has shown PTH (1-34), PTHrP (1-36), and ABL exert time and dose-dependent differential responses in the osteoblast [13]. Many studies have examined PTH signaling in the osteoblast/osteocyte [9, 13, 21, 22, 23, 24, 25, 26], ours is the first to compare these peptides and show that they differentially modulate the osteoblast transcriptome in primary osteoblasts.

RNA-Sequencing data revealed PTH (1-34) regulated the most genes (367), followed by ABL (179 genes) and then by PTHrP (1-36) (116 genes), with PTH (1-34) generally having the greatest fold effects. Gene ontology analyses show biological processes are similar between the three peptides but have some pathway-specific differences. All 3 peptides regulated genes involved in ossification, cAMP signaling and epithelial cell proliferation, while PTH (1-34) and PTHrP (1-36), but not ABL, regulated genes of branching morphogenesis by GO. The latter may reflect the role of PTHrP in mammary gland development [27]. PTH (1-34) and ABL show an almost identical pattern otherwise, including Wnt signaling, hormone transport and secretion, bone mineralization, and actin filament bundle organization. PTHrP (1-36) regulates these pathways differently when compared with either of these peptides. These similarities in pathway expression among PTH (1-34) and ABL may explain why they have similar effects *in vivo* and why PTHrP (1-36), despite being an analog of ABL, did not have the same anabolic effects on bone mineral density [28]. With respect to molecular functions and cell components, PTH (1-34) gave the greatest and most significant effects, while all three peptides regulated phosphoric ester hydrolase activity (a feedback control for cAMP action), G-protein-coupled peptide receptor activity, nuclear and neuropeptide receptor activity, many of which were reflected in the cell component designation of receptor complex, which was significant for PTH (1-34) and ABL.

Based on a close examination of the RNA-Sequencing data we selected a handful of genes that were significantly regulated by these peptides with either similar or differential expression patterns for confirmation via qRT-PCR. Additional mouse osteoblast samples were harvested for subsequent RNA isolation and additional analyses. From these data we were able to confirm several notable expression patterns elicited by these peptides in both the RNA-sequencing data and qPCR.

The first pattern of note which we found in both the qPCR and RNA-Sequencing data is that *Vdr, Cited1, Pde10a, Wnt11*, and *Sfrp4* followed the reported pattern of expression profiling of *Rankl* in response to these peptides: PTHrP (1-36) and ABL result in a moderate increase in gene expression compared with control with PTHrP (1-36) being, in most cases, significantly lower than ABL, and PTH (1-34) producing the greatest increase in expression of these genes compared with control (Fig 5).

VDR is an important factor in serum calcium homeostasis and with 1,25(OH)_2_ vitamin D_3_, is known to bind regulatory elements in the PTH gene promoter to block PTH transcription [29, 30]. A previous report showed that daily administration of PTH (1-34) for 48 h decreased renal *Vdr* mRNA expression by 15% in wild-type mice and determined that intermittent PTH (1-34) is a potent down-regulator of *Vdr* mRNA *in vivo* [31]. However, our data show that *Vdr* mRNA is increased by PTH (1-34) in osteoblasts and this may be explained by the periodicity of treatment and cell type. In fact, one group showed that the PTH1R was present in mandibular/alveolar mouse bone at the earliest stages examined in embryogenesis while the VDR only appeared later with maximal expression at E18 implying it may be induced by PTH or PTHrP action [32]. In a prehypertrophic chondrocyte cell line, PTHrP was shown to substantially increase *Vdr* expression and the authors suggested there was a functional paracrine feedback loop modulating chondrocyte differentiation [33]. In our case, the highest regulation by PTH (1-34) in osteoblastic cells intimates that this hormone may control 1,25(OH)_2_ vitamin D_3_ action on these cells.

The second gene showing the same pattern of regulation as *Rankl, Cited1*, has also been reported to be up-regulated significantly by PTH (1-34); Wt9 osteoblastic cells showed maximal upregulation of *Cited1* after 4 h of PTH treatment which was blocked by PKA inhibition [26]. They also observed that in calvarial osteoblasts derived from *Cited1* knockout mice treated intermittently with a cAMP-selective analog of PTH, [G1,R19]hPTH (1-28), there were greater increases in mineralization, which identifies CITED1 as a negative regulator of osteoblast differentiation. The protein is a transcriptional coactivator of the CBP/p300-mediated transcription complex which interacts with Smads. Its upregulation by PTH (1-34) may be a feedback control.

The general function of phosphodiesterases is to hydrolyze cyclic AMP and cyclic GMP second messenger molecules, thus regulating them as second messengers [34]. PDE10A has the highest affinity for cAMP and most of our knowledge on it comes from studies in the brain [34, 35]. However, a recent study examined *PDE10A* expression in bone marrow-derived mesenchymal stromal cells isolated from a patient cohort undergoing hip replacement therapy [34]. They report that *PDE10A* is upregulated in response to mechanotransduction and that its upregulation impairs osteogenic signals and that an increase in cAMP was the key driver in the observed results. Our data show that *Pde10a* mRNA is upregulated significantly by all three peptides but with PTH (1-34) showing by far the largest increase in its expression. It is most likely that this is feedback regulation to control cAMP concentrations and the highest induction by PTH (1-34) reflects the highest cAMP produced by this peptide in these cells.

SFRP4 has been shown to have deleterious effects on osteoblasts/osteocytes. SFRPs are well known antagonists of Wnt family signaling by directly binding the Wnts as decoy receptors or by forming nonfunctional Wnt complexes via Frizzled (Fz) proteins [36]. Canonical Wnt signaling typically promotes bone formation so any complex that removes or inhibits its function is likely to negatively affect osteoanabolic processes as shown in a study which utilized transgenic mice overexpressing SFRP4 and found that they exhibited comparatively lower bone mass [37]. The study concluded that this deleterious effect could be mostly attributed to a substantial suppression in bone proliferation/formation. Our data show that *Sfrp4* also has the highest upregulation in response to PTH (1-34) when compared to ABL and PTHrP (1-36) and may also be a feedback control mechanism in response to cAMP levels. The greater effect of PTH (1-34) may mean greater restraint on Wnt action in the osteoblast compared to ABL.

Our pathway analyses and subsequent qPCR show that *Wnt 4, Wnt 7b,* and *Wnt 11* all have significantly increased mRNA expression in response to all three peptide treatments compared to control but do not do so with the same pattern. *Wnt11* mimics the *Rankl* expression profile with PTH (1-34) having by far the greatest effect compared with ABL and PTHrP (1-36) while *Wnt4* / *Wnt7b* expression was highest with PTHrP (1-36) and PTH (1-34) having a lesser effect. WNT4 was known to be a non-canonical WNT family member but has been shown to act by both canonical and non-canonical pathways and to have a number of roles in bone [38]. WNT11 appears to stimulate osteogenesis through the canonical pathway and WNT7B seems to also act through the canonical pathway to regulate limb development [39, 40].

The effects of PTH (1-34) and its analogs on the osteoblast transcriptome illuminate the exquisite nature of bone modeling and remodeling. While several hundred genes are regulated by each peptide, we determined that pathway analysis would provide a more systematic way of analyzing the regulatory trends. Perhaps choosing an earlier time point would yield different results, particularly if performed to capture immediate early genes. Nonetheless, *Rankl, Vdr, Cited1, Pde10a, Sfrp4*, and *Wnt11*, which were differentially regulated in a similar pattern by the three peptides suggested that these are affected by the initial differences in cAMP/PKA signaling and may also involve the SIK/CRTC/bZIP pathway, which regulates *Rankl* transcription [13]. Conversely, it is possible that the genes that were similarly regulated by all three peptides are governed by the SIK/HDAC pathway as shown by the similar regulation of *Sost* and *Mmp13*, both of which are controlled through the SIK/HDAC arm. qRT-PCR analysis of these genes in calvarial osteoblasts treated with 8-bromo-cAMP, myr-PKI, si-*Sik*s, si-*Crtc*s, or si-*Hdac*s would be able to confirm this hypothesis [13].

To this end *Vdr, Cited1, Wnt4, Wnt7b, Wnt11, Sfrp4, Epha3,* and *Sost* mRNAs were examined in cells where SIK1/2/3 and CRTC1/2/3 knockdowns were performed followed by treatment with and without PTH (1-34) (Fig 7). The results demonstrated that while knockdowns did not significantly change the mRNA levels of these genes in the basal state, there was substantial regulation of these genes after the knockdowns with PTH (1-34) treatment. *Vdr* mRNA levels were significantly increased, c.f. PTH (1-34) treatment, in response to si-*Sik1* and si-*Sik3* and decreased with CRTC2/3 knockdown similar to our observations with *Rankl* [13 and unpublished data]. *Wnt4* mRNA levels were also similarly affected by SIK1 and CRTC2/3 knockdown, suggesting that these three genes (*Vdr, Rankl, Wnt4*) are regulated through the same pathway.

*Sost* and *Sfrp4* (both WNT pathway inhibitors) seem to be controlled similarly, by SIKs but not CRTCs, possibly both through the Type II HDAC arm, although this would need to be proven for *Sfrp4*. Notably, PTH (1-34) decreases *Sost* expression, which has been thought to be a major part of its anabolic effects on bone, while it increases *Sfrp4* expression, which seems to be a feedback control of the WNT pathway.

PTH (1-34) stimulation of *Wnt7b* and *Wnt11* mRNA levels was significantly decreased by both SIK1/2 and CRTC2 knockdown. This suggests a positive role for SIK action in *Wnt7b* and *Wnt11* transcription. A similar observation has been made for TGF-β-stimulated transcription of *PAI-1* [41] and the authors speculated that the SIKs were involved with required phosphorylation of co-activators binding to P-Smads in the nucleus, but this is an open question, as it is for how SIKs might regulate *Wnt7b* and *Wnt11* transcription by osteoblasts.

It is notable that *Cited1* transcription is not regulated by the SIKs, and this was observed previously, indicating that there are a number of genes that PTH controls independent of this pathway [21]. It is possible that these are directly regulated by PKA-phosphorylation of CREB and do not involve the CRTCs or by some other unknown pathway.

Future research could explore these potential mechanisms by employing advanced experimental approaches such as proteomics, transcriptomics, and systems biology to uncover novel interactions, regulatory elements, or context-dependent effects. Additionally, studying the spatiotemporal dynamics of these proteins during development or tissue homeostasis could provide further insights into their functions and interactions.

The findings in this study highlight the complexity of the genetic and functional events that are triggered by PTH(1-34) and its analogs. Although we discovered many genes that seemingly fit into known paradigms there were many genes that should be further evaluated to understand the importance of PTHR1 and how it affects downstream signaling. A closer examination of some of these genes might reveal intricacies in the interactions of PTHR1 and PTH-derived treatments just as further delineation of which events are attributable to signaling mechanisms triggered by PTH(1-34), PTHrP (1-36), and ABL would allow for further refining of future treatments for osteoporosis.

